# *SaVor* - A Reproducible Structural Variant Calling and Benchmarking Platform from Short-Read Data

**DOI:** 10.1101/2025.11.26.690832

**Authors:** Trevor Mugoya, Arun Sethuraman

## Abstract

Structural variations (SVs) are differences in genomic regions that are larger than 1 kilobase-pair (Kbp) between individuals, and can arise from errant DNA repair mechanisms, whole genome duplications, and transposable element activity across the genome. Recent advances, optimizations, and cost reductions in next generation sequencing technologies have facilitated the exponential increase in the amount of available short read genomic data. Here we present *SaVor*, a flexible, reproducible SV calling workflow that accepts single or multi-lane short-read paired-end Illumina sequence data, or BAM files as input to generate a consensus SV call-set based on user-provided merge parameters. We tested *SaVor* on 1,165 *Arabidopsis thaliana* whole genome sequences and benchmarked its performance on a set of SVs derived from the same accessions using *Lumpy*. Intersection calls i.e. SVs supported by 3 SV callers showed the highest precision (>0.91) while union calls supported by at least 1 caller showed the highest recall (>0.88). We found that the former suffers from decreased recall (<0.51) and the latter decreased precision (<0.57). Depending on the merge strategy, trade-offs in recall and precision need to be considered for downstream analyses of SV call-sets from short-read data. *SaVor* is an open-source Snakemake pipeline and is available on GitHub at https://github.com/ChabbyTMD/SaVor

## Introduction

Structural variants (SVs) are characterized by genomic DNA regions larger than 1 Kbp that exhibit changes in copy number (insertion, deletions or duplications), orientation (inversions), or chromosomal location (translocations) among individuals (Escaramís et al., 2015). Early studies of structural variation, especially in plants were limited by the lack of high-quality reference genomes and the occurrence of polyploidy, which complicates downstream analyses (Yuan et al., 2021). Advances in modern sequencing technologies have led to the availability of numerous high quality plant genomes and improved SV detection capacity (Schwacke et al., 2025). Additionally, improvements in the resolution of small-scale SVs and copy number variants (CNVs) from short-read sequencing data previously unattainable with earlier SV detection techniques such as array comparative genomic hybridization (aCGH) and single nucleotide polymorphism (SNP) arrays (Escaramís et al., 2015). Detecting SVs from short-read sequence data continues to be challenging. Modern short-read sequencing platforms (e.g. Illumina NextSeq^Ⓡ^ and MiSeq^Ⓡ^) generate reads in the range of 150-300 bp (Slatko et al., 2018). This read length is not suitable for ‘direct’ SV detection. Thus, several methods have been developed to detect SVs from short-read mapping signatures. These methods include Read Depth, Read Pair, and Split Read Analyses (Alkan et al., 2011). Read Depth (RD) analysis can detect deletions, duplications, and determine absolute copy number by uncovering discrepancies in read depth distribution across mapped regions. Read Pair (RP) analysis characterizes SV breakpoints for insertions, deletions and inversions by evaluating insert length distributions of mapped read pairs. Split Read (SR) analysis probes read sequences for discontinuities that are signatures of structural variant breakpoints for inversions, translocations, insertions and deletions.

Bioinformatics tools have been developed to detect SVs using one (*CNVnator* (Abyzov et al., 2011)) or a combination of multiple methods such as *Lumpy* (Layer et al., 2014), *Delly* (Rausch et al., 2012), and *Wham* (Kronenberg et al., 2015).

A few challenges arise with SV detection, however, especially in analyses that rely on a single tool/method. First, utilizing a single tool or method has been reported to have high observed false discovery rates and wide variation in sensitivity of SV detection (Sedlazeck et al., 2018). Second, tools based only on RD methods are orientation agnostic and thus can only detect unbalanced SVs such as deletions and duplications (Alkan et al., 2011). Third, RP analysis fails to distinguish between tandem duplications and novel insertions due to their similar mapping pattern (Mahmoud et al., 2019).

To address these challenges, SV calling tools have been developed to evaluate SV breakpoint evidence from all, or a combination of RD, RP, and SR methods to generate SV catalogs (Cameron et al., 2019). Alternatively, multiple bioinformatics tools can be used in a single analysis workflow to generate an ensemble SV call-set that represents a consensus of some, or all SV callers (Ho et al., 2020). Irrespective of the method employed, the implementation of one or multiple tools can be best achieved by a workflow management system e.g. Snakemake or Nextflow (Di Tommaso et al., 2017; Köster & Rahmann, 2012). While individual scripts tend to be non-portable and non-reproducible, workflow management languages have built-in support for package managers like *conda* that allow easy replicability of computational environments. Snakemake facilitates this by building *conda* environments based on environment YAML file(s) detailed by the developer that are dynamically activated and called depending on the rule being executed (Köster & Rahmann, 2012).

Here we present *SaVor*, a reproducible pipeline written in Snakemake that applies an ensemble of three short-read based SV callers (*Lumpy*, *Wham* and *Delly*) and subsequently generates a consensus set of SVs (Kronenberg et al., 2015; Layer et al., 2014; Rausch et al., 2012). *SaVor* bundles a configurable SV benchmarking submodule that provides SV F1, precision and recall scores for duplications, deletions, and inversion structural variant types generated by *Truvari* (English et al., 2022).

### SaVor Overview

*SaVor* is an end-to-end structural variant calling workflow that takes either aligned or unaligned short-read sequence data as input. If aligned data are provided, the inputs are reference-mapped, coordinate-sorted, and de-duplicated Binary Alignment Map (BAM) files, in addition to the reference genome. If unaligned data are provided, the supported inputs are paired-end FASTQ files from one or multiple sequencing lanes and a reference genome (FASTA). A single Variant Call Format (VCF) file containing a union of at least one, or an intersection of all structural variant caller outputs merged across all samples is the final output of the pipeline. Three structural variant callers, *Lumpy*, *Delly* and *Wham* are integrated into *SaVor*, each utilizing the same read alignment, reference genome, and its associated indices to independently call SVs. Raw per-sample SV calls generated by each tool are then merged by SURVIVOR according to a user-specified option in the *sv_merge* directive in the workflow *config.yaml* file for the number of callers supporting an SV (Jeffares et al., 2017). Each sample VCF is then filtered using *bcftools* (ver. 1.22) (Danecek et al., 2021) for SVs greater than 10,000 base pairs in length (Danecek et al., 2021). All sample VCFs are then merged into a single consensus VCF file using SURVIVOR.

An optional benchmark submodule is also implemented in the *SaVor* workflow. When enabled, the consensus VCF file is used as input to generate three comparison VCF files for deletions, inversions, and duplications SV types for benchmarking. “Ground truth” VCF files provided by the user in addition to the comparison VCF files generated by the module for each SV type are passed to the benchmarking tool *Truvari* (English et al., 2022) to generate F1, recall, and precision metrics (Fig 1). Snakemake’s native support for *conda* allows *SaVor* to automatically deploy discrete *conda* environments depending on the currently active job(s). *SaVor* uses tools that are distributed via the *bioconda* (Grüning et al., 2018) and *conda-forge* (conda-forge community, 2015) channels. Conda environment recipes are described in YAML files in the “SaVor/envs/” directory. The entire *SaVor* workflow is managed in a single configuration “config.yaml” file, with additional directives to specify the SV call merging approach, location of ground-truth VCF files and to activate *SaVors*’ benchmark module.

**Figure 1.**
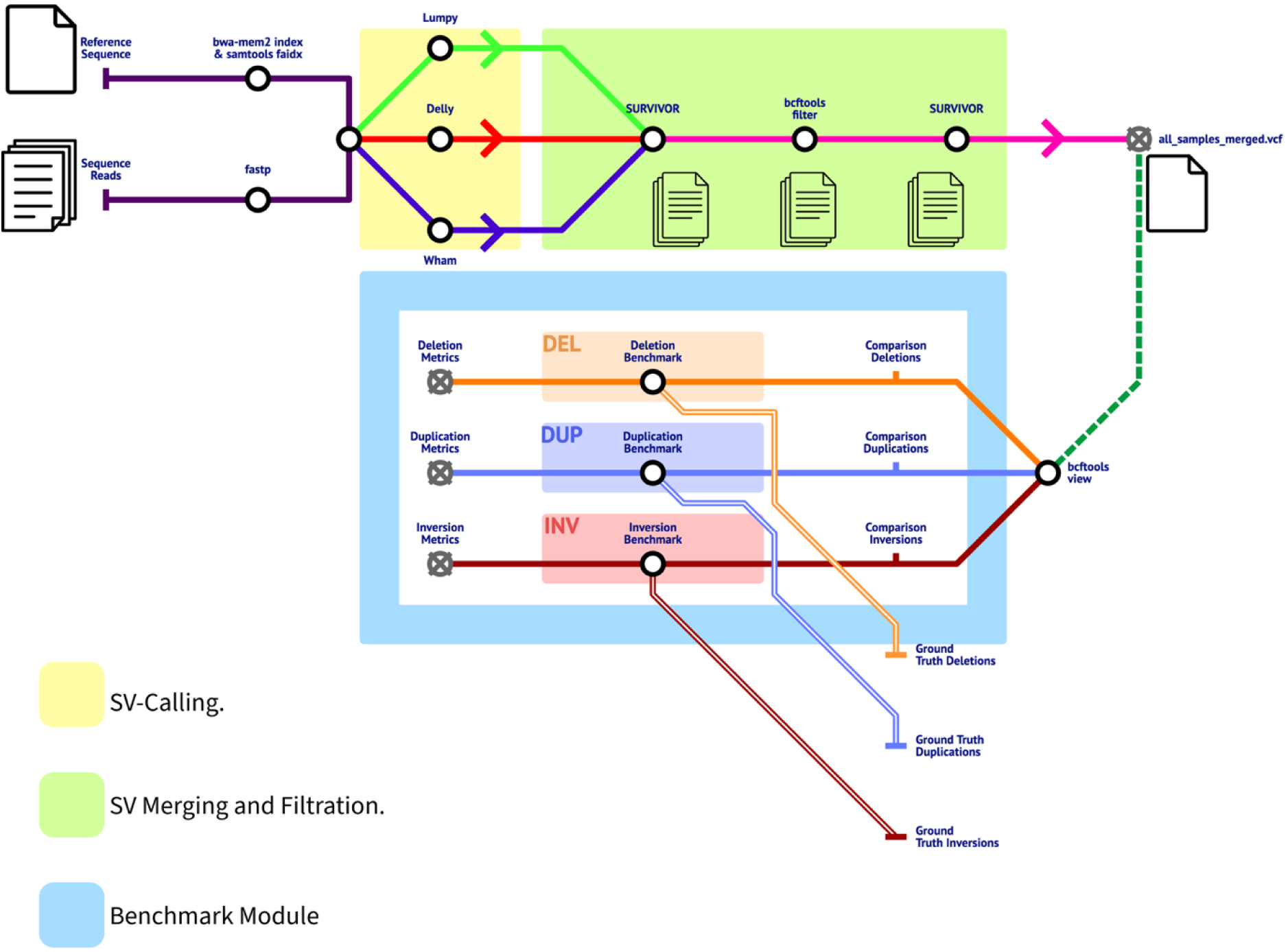
SaVor overview. SaVor is an end-to-end structural variant calling module written in Snakemake (Köster & Rahmann, 2012). SaVor accepts paired-end FASTQ reads and a suitable reference genome(FASTA) to generate read alignment files (BAM) and their associated indices (BAI) that are passed on to the SV calling tools Lumpy, Wham and Delly. A merged SV call-set VCF files is the final output of the main workflow. Savor’s benchmark module, when activated through the “svBenchmark” directive in the workflow “config.yaml”, takes the merged call-set evaluates deletion, duplication and inversion calls against a user-provided structural variant ground-truth call-set and outputs precision, F1 and recall metrics.

## Results

With the exception of *Lumpy,* which represents the “best-case” scenario of perfect SV call matching in our tests, across all considered SV types, *SaVors’* union merging option (*sv_merge=1*) improved recall scores by 13.1%, 9.57% and 87.29% for duplications (0.96), deletions (0.88), and inversions (0.89) respectively compared to the second-best single SV caller, *Delly*. The SV calls generated by the intersection approach (*sv_merge=3*) decreased recall for all SV types with scores of 0.51, 0.59, and 0.07 for duplications, deletions, and inversions respectively. We observed precision scores for the union SV calls being lower for duplication and deletion SV types in comparison to singular callers. The precision values were 0.43, 0.53, and 0.23 for duplications, deletions, and inversions respectively. Inversions were a notable exception, showing a 53% improvement in precision score over *Delly*. However, the intersection SV calls showed a significant improvement over the union SV calls; *Wham* and *Delly* in precision with scores of 0.98, 0.91 and 0.94 for duplications, deletions and inversions respectively. *SaVor’s* intersection approach showed marginal improvements in F1 score over the next best caller, *Delly*, with duplications and deletions with scores of 0.66 and 0.71. Interestingly, the intersection SV calls had the lowest F1 scores for inversions at 0.135, while *SaVor’s* union SV calls had the second highest at 0.37.

We observed that *SaVor’s* intersection merging approach yielded the lowest false discovery rate (FDR) of all considered SV types. *Delly* had the highest FDR for inversions at 0.849 followed by *SaVor’s* union SV calls at 0.765. *Wham* performed the second best of the individual structural variant callers with FDRs of 0.268, 0.482 and 0.688 for deletions, duplications, and inversions respectively (Fig 3).

## Discussion

*SaVor* is a reproducible, flexible structural variant calling and benchmarking workflow that can be easily configured by the user to run on either local Linux machines or high-performance compute (HPC) clusters with the SLURM scheduler.

All structural variant calling tools in *SaVor* are implemented as discrete Snakemake rules that are subsequently executed in compartmentalized *conda* environments that are dynamically activated based on the current tool Snakemake is running. This facilitates reproducibility of computation environments, mitigates dependency conflicts between tools, and allows for the flexibility to substitute different tools in separate Snakemake rule files.

We tested *SaVor* with 1,165 publicly available short-read whole genome sequences of *Arabidopsis thaliana* samples from a global distribution derived from BioProjects PRJNA273563 and PRJNA293798 (The 1001 Genomes Consortium, 2016). In total, 3.9TB of raw FASTQ files were downloaded from the NCBI SRA and the workflow was executed on five 16 core Intel^®^ Xeon^®^ E5-2620 CPU, 128 GB RAM nodes with computational jobs distributed to the SLURM scheduler via the Snakemake “cluster-generic” executor plugin (*Snakemake-Executor-Plugin-Cluster-Generic*, n.d.). The *SaVor* workflow generated a union SV call-set containing 540,023 records and an intersection SV call-set containing 36,219 records. Total execution time for the entire workflow was approximately 150 wall-clock hours for both runs.

Our evaluation metrics highlight that depending on the merging approach used, there is a trade-off between precision or recall of the final SV call-set. Opting for an intersection approach will improve precision at the cost of recall and a smaller set of final variants generated by the workflow. The benefit of this approach is seen in FDR, which is lower than two of its constituent SV callers. This implies that the intersection approach greatly reduces the number of false positive SV calls in the generated SV call-set. A downside of this approach, however, is the high false negative rate, where SV calls present in the ground truth set were not matched to calls in the comparison set. This may indicate that the intersection approach is highly stringent on calls made by one or two callers, potentially not capturing biologically relevant SVs. On the other hand, the union calling approach yields the highest recall at the cost of precision. The benefit of this approach is that one can conserve most of the SV calls made by all three of the SV callers implemented in SaVor. However, if a set of SVs can be uniquely detected by one or two tools due to the functionality of their respective algorithms, their inclusion in the final consensus may be flagged as false positive hits during benchmarking. This can be seen in Fig. 2, where a large number of false positives contributes to decreased precision and F1 score. This consequently inflates the FDR of the union SV calls, and this is indirectly reflected in the FDRs of the individual SV callers *Wham* and *Delly*. Therefore, there could be unique biologically relevant calls predicted by either of these tools that are being flagged as false positives as they do not match calls in the ground truth set.

**Figure 2.**
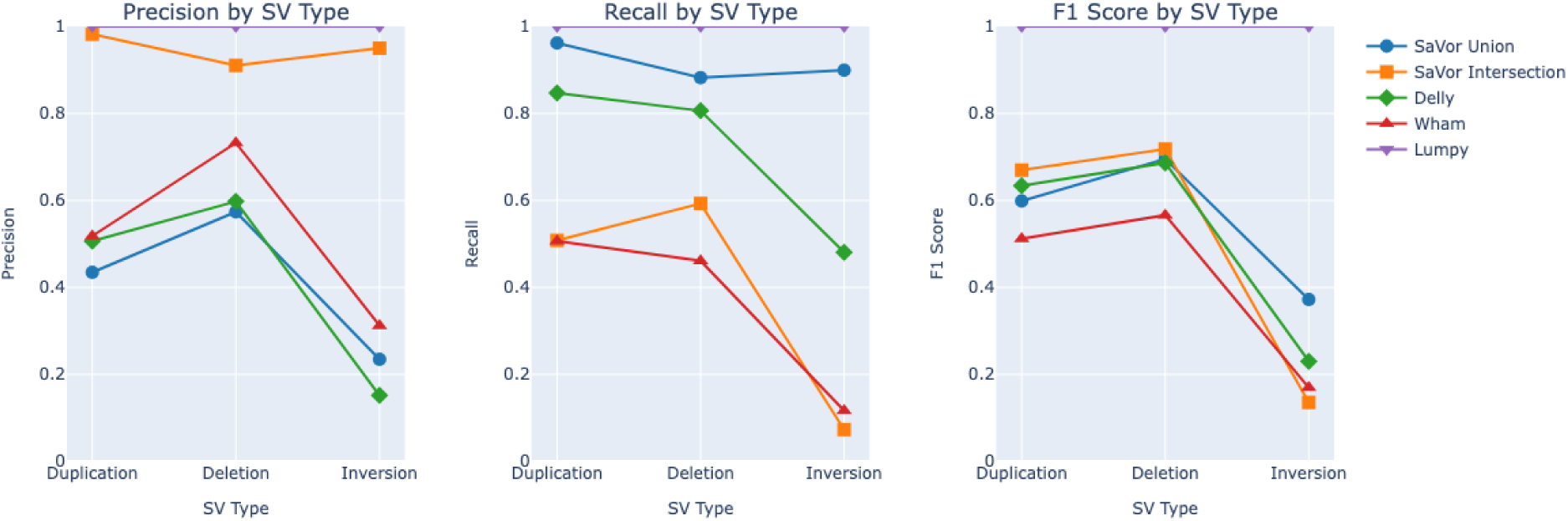
SaVor evaluation metrics (precision, recall and F1) against the individual variant callers implemented within the workflow.

**Figure 3:**
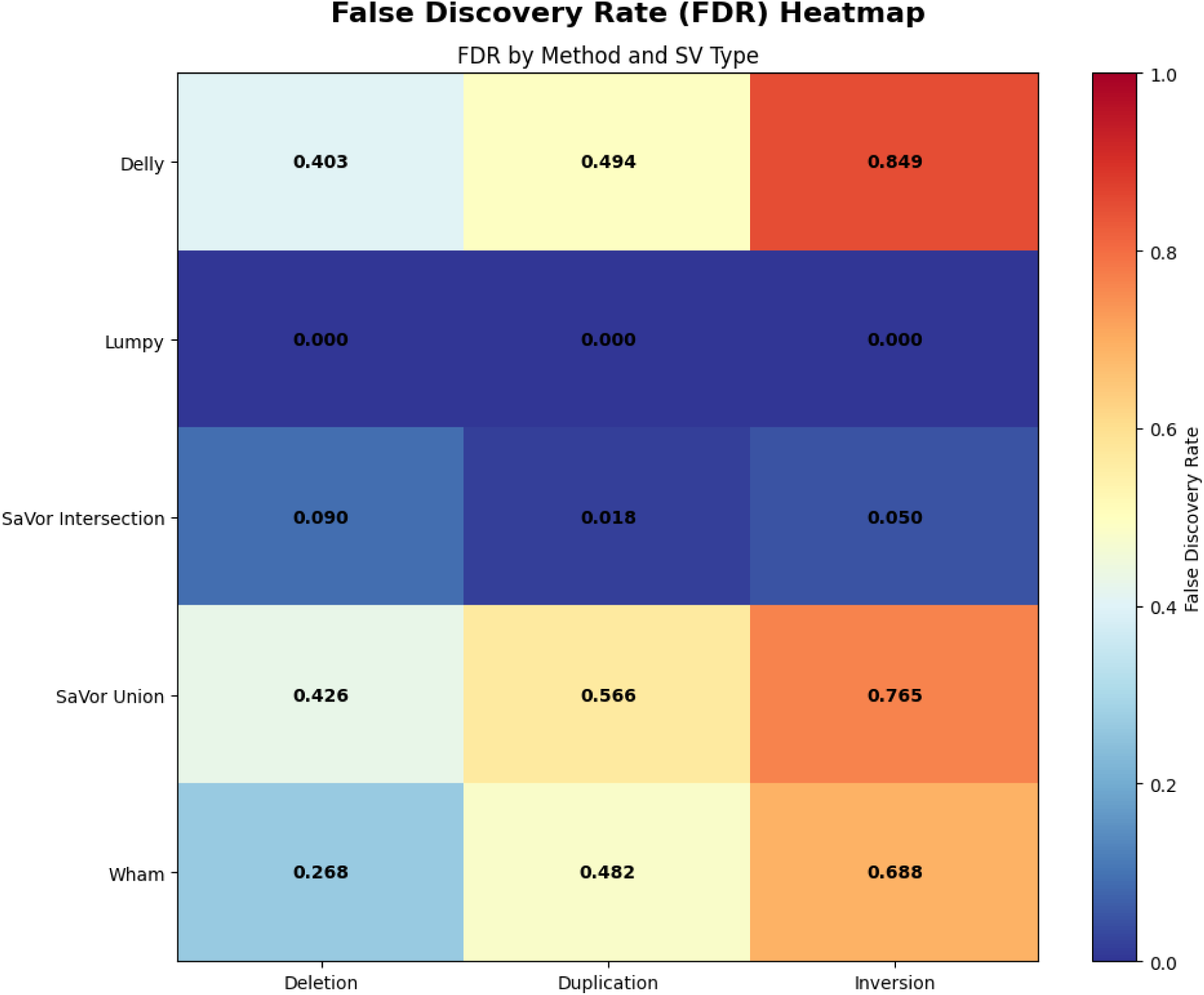
False Discovery Rate Heatmap of SaVor Union and Intersection benchmark results alongside individual SV callers.

An additional limitation of the experimental setup in this study was the lack of a curated modern NGS structural variant call-set. The 1001 genome initiative generated an SV call-set on 80 *A. thaliana* accessions with the SHORE pipeline in a non-VCF format that is not compatible with the benchmark tool implemented in this workflow (Cao et al., 2011). We therefore contend that with more accurate and precise SV call-sets generated via long-read sequencing, *SaVor* would serve as a fast, reliable, and reproducible SV-calling pipeline for short-read resequencing projects. Further development to improve on the *SaVor* workflow will add ease of use features, such as improved SV tool parameters via a unified configuration file, and additional rules to ignore low-complexity and repeat regions to avoid ambiguous or erroneous SV calls regions.

## Methods

### Experimental Setup

The *SaVor* workflow was tested on 1,165 paired-end whole genome short-read sequences from a global distribution of *Arabidopsis thaliana*. The data were obtained from NCBI SRA BioProjects PRJNA273563 and PRJNA293798 (The 1001 Genomes Consortium, 2016; Zou et al., 2017). A list of accessions for all samples and their respective provenances is available in Supplement File 1. Reads were mapped to the *Arabidopsis thaliana* TAIR10 reference assembly (GCF_000001735.3).

### Empirical Data Preprocessing

Raw FASTQ files for the selected *A. thaliana* accessions were downloaded using the *ffq* tool implemented in the *get_fastq_pe* rule (Gálvez-Merchán et al., 2023). Quality control and read trimming was performed by *fastp* (ver. 0.20.1) using default parameters (Chen et al., 2018). The *A. thaliana* TAIR10 (GCF000001735.3) reference was indexed by *bwa-mem2 index* (ver. 2.2.1) and *samtools faidx* (ver. 1.14) (Danecek et al., 2021; Vasimuddin et al., 2019). Subsequent reference-based mapping and sorting was performed with *bwa-mem2* and *samtools merge* respectively (Danecek et al., 2021; Vasimuddin et al., 2019). Each sample BAM file was marked for duplicates by *sambamba* (ver. 0.8.0) with default parameters to generate final BAM files (Tarasov et al., 2015). The BAM files in addition to the TAIR10 reference genome and their associated indices were passed in as input to the structural variant calling.

### Structural Variant Calling

Three popular structural variant callers, *Lumpy, Delly* and *Wham* (Kronenberg et al., 2015; Layer et al., 2014; Rausch et al., 2012), were each implemented in their own ‘snakefiles’ within the *SaVor* workflow. All three variant callers are compatible with, and require bwa-mapped, coordinate-sorted, and deduplicated alignments as input to call structural variants. *Lumpy* (ver.0.3.1) is based on a general probabilistic framework that can flexibly consider multiple lines of evidence such as paired-end sequence alignments, split read mappings, breakpoint probability distributions, and known SV genomic coordinates, if “ground truth” sets are available (Layer et al., 2014). Discordant and split read alignments were extracted from samples final BAM file input using *samtools view -F* with the bitwise flag 1294 (extract properly paired reads (0x2), in which the read or its mate is unmapped (0x4), (0x8), the read is not primary alignment (0x100) and the read is a PCR or optical duplicate (0x400); (Quinlan, 2012/2025; *SAM Format Flag*, n.d.) and a custom *Lumpy* helper script, *extractSplitReads_BwaMem*, respectively. Extracted alignments were subsequently sorted with *samtools* prior to SV calling with the main *Lumpy* executable (Danecek et al., 2021). Contig IDs and lengths were then added to *Lumpy* generated SV VCF file headers with the *bcftools reheader* command in the *fix_lumpycall_header* rule (Danecek et al., 2021).

Delly (ver.1.1.6) analyzes mapping patterns of short and long-range sequence read pairs in addition to split read analysis to refine structural variant breakpoint resolution to the single nucleotide scale (Rausch et al., 2012). Coordinate-sorted and deduplicated *bwa-mem2* alignments were used as inputs to Delly implemented in the *delly_call* rule to call SVs and subsequently output per sample Binary Call Format (BCF) files. For consistency with downstream tools, the BCF SV output files from Delly were converted to VCF files by the *delly_vcf* rule (Danecek et al., 2021).

Wham (Whole-genome Alignment Metrics) (ver.1.8.0) detects structural variants using a combination of “mate-pair mapping, split read mapping, soft-clipping, alternative alignment and consensus sequence-based evidence” to refine breakpoint resolution (Kronenberg et al., 2015). Coordinate sorted and deduplicated bwa-aligned BAM files, in addition to a CSV file containing a list of reference contig/chromosome IDs are used as inputs to *Wham* implemented in the *wham_call* rule.

### Post-Processing

Output VCF files from all three structural variant callers were used as inputs to the SV post-processing rule file. First, each per sample VCF file was sorted with *bcftools* followed by call-merging using SURVIVOR (Jeffares et al., 2017). To consolidate all SVs called in a sample into one VCF file, we employed two sets of merge parameters to SURVIVOR. One to generate a union SV call-set among all callers (1000 1 1 1 0 50) and another that would generate an intersection set with SV calls supported by all three SV callers (1000 3 1 1 0 50). The common parameters between the two sets ensure SV calls were merged if the maximum distance between breakpoints was less than 1000 bp, SV type and strand match, and minimum SV size was greater than 50 bp. The latter set of parameters (1000 3 1 1 0 50) is *SaVor’s* default mode. However, the number of callers needed to support a particular SV call for merging purposes can be modified by the user in the workflow “config.yaml” file by changing the value of the *sv_merge* directive. The unified sample SV VCF files were then further filtered to remove structural variants larger than 10Kbp with *bcftools* (Danecek et al., 2021). All per sample variant calls were then merged across the outputs of the structural variant callers using SURVIVOR (Jeffares et al., 2017). Additional processing to ensure the symbolic allele in the ALT field of the VCF file matched the records’ structural variant type was performed using a custom *awk* script implemented in the *post_merge_process* rule. From this merged call-set, deletions, inversions, and duplications were extracted into their own separate files generated by the benchmarking submodule.

### Benchmarking

SV calls using the same *A. thaliana* accessions were generated by *Lumpy* and genotyped by *Svtyper* (ver.0.7.0) (Chiang et al., 2015) were used as ground truth sets to evaluate the recall, F1, and precision of the SaVor final consensus SV call-set. In total 43,701 deletions, 11,018 duplications and 26,321 inversions respectively were present in the base VCF files. The benchmark submodule within *SaVor* split the final merged VCF output file of the main workflow into three separate VCF files using *bcftools*, each containing a singular SV type (i.e. Deletions, Duplications and Inversions) as a “comparison” set. The structural variant comparison and analysis toolkit, *Truvari* (English et al., 2022), was used to generate F1-score, precision, recall, and genotype concordance metrics for each SV type. Structural variant sequence (-p) and size similarity (-P) parameters were both set to 50%. Reference distance, the maximum allowed distance within which a comparison call’s start and end position must reside in relation to a “ground truth” calls’ start and end position in the reference genome to qualify as a match, was set to 2000bp. SV type matching (True), minimum reciprocal overlap (0%), maximum break-end distance (100bp), and other *Truvari* parameters were left as default. Each SV type VCF file was first sorted and indexed with *bcftools* and *tabix*, respectively, prior to benchmarking with the “truvari bench” command. *Truvari* metrics are computed as follows.

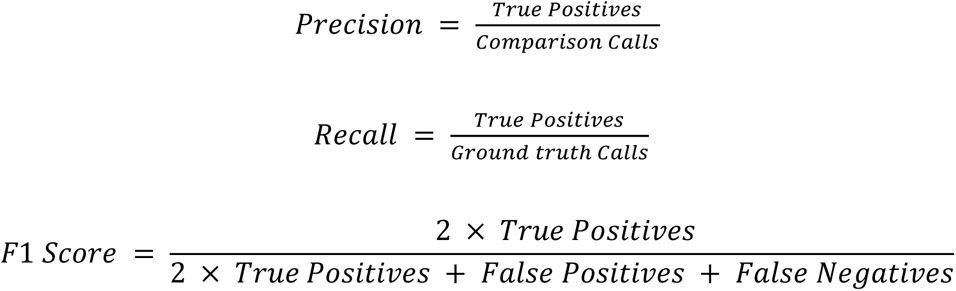

Where “True Positives” are the number of calls in the comparison set that pass all matching criteria to a call in the “ground truth” set, “False positives” are the number of calls in the comparison set that did not match any call in the “ground truth” set and “False Negatives” are the calls in the “ground truth” set that had no corresponding matches in the comparison set.

Additionally, false discovery rate (FDR) was calculated from the number of true positive and false positive calls reported by *Truvari* as follows.

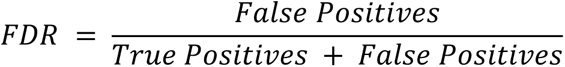

## Acknowledgements

This work was supported by NSF CAREER 2147812, NIH 1R15GM143700-01, and a 2022 CSUBIOTECH Research and Development to PI Arun Sethuraman. All computations described here were performed on the *mesxuuyan* and *anthill* HPC’s at San Diego State University which were funded by NSF ABI 1564659 to PI Arun Sethuraman and co-PI Jody Hey (Temple University), and startup funds to Sethuraman. Trevor Mugoya was supported by the 2023 SDSU Presidential Graduate Research Fellowship, 2023 SDSU Masters Fellowship, and DE-SC0025673 to PI Archana Anand (San Francisco State University) and co-PI Arun Sethuraman. We would like to thank members of the Sethuraman Lab for comments and suggestions on early versions of *SaVor*. We acknowledge the invaluable inputs from Tim Sackton and Cade Mirchandani throughout the development of *SaVor*.

## Data Availability

All genomic data accessions from the *Arabidopsis thaliana* datasets utilized in this study are provided in the Supplementary Material (NCBI SRA BioProjects PRJNA273563 and PRJNA293798). SaVor is a Snakemake pipeline and is available on GitHub:

https://github.com/ChabbyTMD/SaVor

## Author Contributions

AS conceived the study, obtained funding, directed the research, and co-wrote the manuscript. TM designed and wrote the pipeline, performed all analyses, and co-wrote the manuscript.

